# Smear positive tuberculosis and genetic diversity of *M. tuberculosis* isolates in individuals visiting health facilities in South Gondar Zone, northwest Ethiopia

**DOI:** 10.1101/616789

**Authors:** Amir Alelign, Beyene Petros, Gobena Ameni

## Abstract

**Background:** Tuberculosis (TB), a bacterial infectious disease, persisted to be a public health concern in many developing countries. However, lack of enough data concerning the public health burden and potential risk factors for the disease hampered control programs in target areas. Therefore, the present study aimed in determining the prevalence of TB and genetic diversity of *M. tuberculosis* isolates in South Gondar Zone, northwest Ethiopia.

**Methods:** A cross-sectonal study was conducte between March 2015 and April 2017. Bacteriological examination, region of difference (RD) 9 based polymerase chain reaction (PCR) and spoligotyping were used to undertake the study.

**Results:** The overall prevalence of smear positive all forms TB was 6.3% (186/2953). Extra pulmonary TB (EPTB) was clinically characterized on about 62.4% (116/186) TB-positive cases. Some of the patients’ demographic characteristics such as [patients’ origin (chi-square (χ^2^) value; 62.8, p<0.001) were found to be significantly associated risk factors for the occurrence of TB in the study area. All the mycobacterial isolates were found to be *M. tuberculosis.* Among the 35 different spoligotype patterns identified, 22 patterns were shared types. The three dominantly identified families were T, CAS and Manu, each consisting of 46.9%, 24.0% and 10.4% of the isolates, respectively

**Conclusion:** The presented study revealed that TB continued to be a public health problem in South Gondar Zone which suggests a need of implementing effective disease control strategies.

## Background

Tuberculosis (TB), an infectious disease caused by members of *Mycobacterium tuberculosis* (*M. tuberculosis*) complex (MTBC) remains to be one of the public health threats in many parts of the world. Despite the tremendous progress in combating TB, the disease still stands as a leading cause of death worldwide [1]. In 2017, an estimated 10.0 million incident cases of TB were reported globally, which is equivalent to 133 cases per 100, 000 population [1]. there were an estimated 1.3 million deaths among HIV-negative people due to the disease, and an additional 300 000 deaths from TB among HIV-positive people. Of which, about 25% of the total cases occurred in the African Region [1].

Currently, Ethiopia is one of the 30 high TB burden countries and also one of the 14 countries with in the three lists of the new WHO classification of high burden countries (high TB, high MDR-TB and high TB/HIV burden countries) [2]. The estimated total TB incidence and HIV-negative TB mortality in Ethiopia in 2016 were 177 and 25/100,000 population, respectively, whch is greater than the global average of 140 and 17/100,000, respectively. [2]. All the above facts suggested that TB is stll a major public health concern in Ethiopia

The emergence of HIV/AIDS coupled with socio-demographic factors and patients’ characteristics such as previous history of TB, place of residency, overcrowding, malnutrition and lack of awarness towards the disease transmission were reported to be some of potential risk factors for the spread of TB in Ethiopia and elsewhere [3-7].

Despite the importance of more data on TB burden, few studies were conducted in Ethiopia and paticularly in the Amhara Region. On top of that, majority of these studies were conducted only on pulmonary TB patients recruited from urban government hospitals or other specific settings such as public prisons [8-10]. Such studies, however, do not provide enough information about the burden of all forms of TB in the general population as well as among patients from rural and semi-urban areas. Therefore, the present study was conducted to determine the prevalence of all forms of TB (pulmonary and extra pulmonary) and its associated risk factors among rural and semi-urban TB suspected residents who were attending health facilities in the study area.

## Materials and methods

### Study area and populaton

The study was conducted in South Gondar Zone, Amhara Region, northwest Ethiopia. Debre Tabor is the capital town of the Zone. It is 666 kms from Addis Ababa and 99 kms from the capital city of Amhara Region, Bahir Dar. There are 10 districts (third administrative unit) in the Zone. South Gondar zone has a total population of 2,051,738, and a population density of 145.56; About 90.47% of the total populaton were rural inhabitants. A total of 468,238 households were counted in this Zone, which results in an average of 4.38 persons to a household and 453,658 housing units [11]. Like the rest of the zones in the northwestern part of the country, the livelihood of the community in South Gondar was largely depended on subsistence agriculture [12].

### Study design

A Cross-sectional study was conducted between March 2015 and April 2017 on smallholder farmers in South Gondar Zone. All forms of smear positive TB patients who visited the district health centers and perpheral rural health posts, zonal and regional referal hospitals (Debretabor, Felegehiwot and Gambi hospitals) were included in this study. TB patients who were under treatment prior to the start of the study and those under 5 years of age were excluded from the study. For convenience, the patient sources were reduced from 10 to 8 districts. Questionnaire survey to assess the socio-demographic and other risk factors for the occurence of TB was also conducted.

### Sample size and sampling techniques

The sample size for the present study was determined at district level using a population proportion with specified absolute precision formula [13].

n = z^2^p (1−p)/d2, where n = the number of TB suspects, z = standard normal distribution value at 95 % CI which is 1.96, p = expected prevalence of tuberculosis among the populaton (0.05, from literature in the study area [3]), d= absolute precision, taken as 5 % (we took a larger d due to resource limitations). Accordingly, the sample size calculated, including a 10% non-responsive rate was 396. Hence, for 8 districts in which the study was conducted, a total sample size of 3168 was determined.

#### Inclusion criteria

All TB suspected smallholder farmers visiting the differnt health facilities in South Gonder Zone, residng in rural and semi-urban settings, who were above 5 years of age, were included in the present study.

### Data collection

#### Sociodemographic data

After eligible subjects were recruited and concented on the study, socio-demographic data of the study participants was collected using a structured questionaire. Data on patient characteristics such as age, sex, district of residency, education status, previous exposure to TB, raw milk consumption habit, and family TB history were collected. Moreover, clinical data on the form of TB cases as either pulmonary or extra pulmonary was included.

#### Sputum sample collection and processing

Paired morning-spot sputum samples from clinically suspected pulmonary TB patients and spot fine needle aspirate (FNA) samples from suspected TB lymphadenitis patients were collected by trained laboratory technicians and pathologists. Sputum samples were collected in standard sterile containers and FNA specimens were collected with cryo-tubes with phosphate buffer saline (PBS) pH 7.2. The sputum and FNA samples were processed for smear microscopic examination as of standard protocols [14]. The samples were first digested and concentrated/decontaminated by the NALC-NaOH method (N-acetyl-L-cysteine-Sodium hydroxide). Smears of the final deposits from the various specimens were stained by the Ziehl-Neelsen (Z-N) method and examined under oil immersion using a binocular light microscope.

#### Mycobacterium culture

The samples were processed for culturing according to the standard methods described earlier [15, 16]. Both sputum and FNA samples were cultured at the Bahir Dar Regional Health Research Laboratory Centre. Briefly, about 100 μl of the sample suspension was inoculated on four sterile Lowenstein Jensen (LJ) medium slopes (two were supplemented with pyruvate and the other two with glycerol) and then incubated at 37°C with weekly examination for growth. Cultures were considered to be negative when no visible growth was detected after eight weeks of incubation. The presence of AFB in positive cultures was detected by smear microscopic exa-mination using the ZN staining method. AFB positive cultures were prepared as 20% glycerol stocks and stored at –80°C as reference. Heat-killed cells of each AFB isolate were prepared by mixing ∼2 loopfuls of cells (≥ 20μl cell pellet) in 200μl distilled H_2_O followed by incubation at 80°C for 45 minutes for the release of DNA after the breaking the cell wall. The heat killed cells were transported to the laboratory at Aklilu Lemma Institute of Pathobiology, Addis Ababa University, for molecular typing.

#### Region of difference (RD) 9-based polymerase chain reaction

To differentiate *M. tuberculosis* from other members of the *M. tuberculosis* complex (MTBC) species, RD9 based PCR was performed according to protocols previously described [17]. *M. tuberculosis* H37Rv, *M. bovis* bacille Calmette-Guérin (BCG) were included as positive controls and water was used as a negative control. Interpretation of the result was based on bands of different sizes (396 base pairs (bp) for *M. tuberculosis* and 375bp for *M. bovis*) as previously described [18].

#### Spoligotyping

Isolates genetically identified as *M. tuberculosis* were spoligotyped for further strain identification. Spoligotyping makes use of the variability of the MTBC chromosomal direct repeat (DR) locus for strain differentiation as previously described [19]. A PCR-based amplification of the DR region of the isolate was performed using oligonucleotide primers derived from the DR sequence. The amplified product was hybridized followed by subsequent membrane washing processes. Known strains of *M. bovis* and *M. tuberculosis* H37Rv were used as positive controls, whereas Qiagen water (Qiagen Company, Germany) was used as a negative control. Hybridized DNA was detected by the enhanced chemiluminescence method. The presence or absence of spacer was used as key for the interpretation the result.

#### Identification of *M. tuberculosis* strains and lineages

A web-based spoligotype database was utilized to assign shared international types (SITs) for the isolates [20]. Strains matching a preexisting pattern in the database were identified with SIT number otherwise considered as orphans. An online TB lineage tool was used to predict the lineages/family of *M. tuberculosis* using knowledge based Bayesian network (KBBN) [21].

### Data analysis

Data were double entered to Excile file format and statistical analysis was performed using SPSS software version 20. Descriptive statistics were used to depict the socio-demographic variables. Chi-square (χ^2^) was used to determine the possible association of risk factors with the occurrence of TB. Odds ratio (OR) was used to show the association of patient characteristics with the clustering rate of mycobacterial isolates. Results were considered statistically significant whenever p-value was less than 5%.

### Ethics approval and consent to participate

Ethical clearance for the study was obtained from the Ethical Committee of Addis Ababa University, Department of Microbial, Cellular and Molecular Biology (Ref. CNSDO/491/07/ 15). In addition, written permission was sought from the Amhara Regional Health Bureau Research Ethical Committee (Ref. HRTT/1/271/07). Each study participant was consented with a written form (S1 Appendix) and agreed to participate in the study after a clear explanation of the study objectives and patient data confidentiality. In case of participants under the age of 18 years, consent was obtained from their parents/guardians.

## Results

### Socio-demographic characteristics of the study participants

A total of 3168 TB-suspected individuals, with a response rate of 93.2% (2,953/3168) participated in the study. Of these, 1,733 (58.7%) were males and 1, 220 (41.3%) were females. Other socio-demographic characteristics such as patient origin, age, education status, status of prior exposure to TB within the family members and their milk consumption habits are presented and their association with TB prevalence has been determined (Table 1).

**Table 1.** Socio-demographic characteristics of study participants and their association with tuberculosis prevalence, South Gondar Zone, northwest Ethiopia (N= 2,953) (2015-2017).

| Characteristics | Number, n (%) | AFB smear result | | $\chi^2$(df) | p-value |
| --- | --- | --- | --- | --- | --- |
|  |  | Positive (%) | Negative (%) |  |  |
| **Patient origin (Districts)** |  |  |  |  |  |
| Dera | 485(16.4) | 48 (9.9) | 437 (90.1) | 62.80 (7) | <0.001 |
| Ebinat | 256(8.7) | 17(6.6) | 239(93.4) |  |  |
| Este | 308(10.4) | 10 (3.2) | 298 (96.8) |  |  |
| Farta | 424(14.4) | 8 (1.9) | 416 (98.1) |  |  |
| Fogera | 494(16.7) | 56 (11.3) | 438 (88.7) |  |  |
| Gayint | 367(12.4) | 9 (2.5) | 358 (91.0) |  |  |
| Libo kemkem | 375(12.7) | 28 (7.5) | 347 (92.5) |  |  |
| Simada | 244(8.3) | 10 (4.1) | 234 (95.9) |  |  |
| **Age group (year)** |  |  |  |  |  |
| <18 | 428(14.5) | 21 (4.9) | 407 (95.1) | 35.46 (4) | <0.001 |
| 18-30 | 676(22.9) | 66 (9.8) | 610 (90.2) |  |  |
| 31-43 | 757(25.6) | 50 (6.6) | 707(93.4) |  |  |
| 44-56 | 486(16.5) | 37 (7.6) | 449 (92.4) |  |  |
| >56 | 606(20.5) | 12 (2.0) | 594 (98.0) |  |  |
| **Sex** |  |  |  |  |  |
| Male | 1,733(58.7) | 100 (5.8) | 1,633 (94.2) | 1.98(1) | 0.159 |
| Female | 1,220(41.3) | 86 (7.0) | 1,134 (93.0) |  |  |
| **Education status** |  |  |  |  |  |
| Illiterate | 1654(56.0) | 117 (7.1) | 1,537 (92.9) | 8.06 (4) | 0.089 |
| Adult education | 225(7.6) | 11 (4.9) | 214 (95.1) |  |  |
| Primary school | 790(26.8) | 37 (4.7) | 753 (95.3) |  |  |
| Secondary school | 169(5.7) | 15 (8.9) | 154 (91.1) |  |  |
| Higher education | 115(3.9) | 6 (5.2) | 109 (94.8) |  |  |
| **TB history in the family** |  |  |  |  |  |
| Yes | 2,187 (74.1) | 134 (6.1) | 714 (93.9) | 182.03(1) | <0.001 |
| No | 766 (25.9) | 52 (6.8) | 2,053 (93.2) |  |  |

N: Total number of suspected cases; n: Number of cases in each category; AFB: Acid-fast bacilli;
$\chi^2$ : Chi-square; df: degree of freedom.

### Prevalence of tuberculosis among the study participants

The overall prevalence of smear positive all forms of TB was 6.3% (186/2953). The study showed variability in TB prevalence among the different socio-demographic characteristics such as sex, age and patient origin (districts). Accordingly, males (3.4% (100/2953), age groups from 18 to 30 years (2.2% (66/2953), and Fogera district (1.8% (56/2953), had the highest prevalence. Moreover, the majority of smear positive cases (72.0%) had history of TB in their family (Table 2).

**Table 2.** Number of smear positive TB cases categorized by sex, patient category and TB type for study participants, South Gondar Zone, northwest Ethiopia (2015-2017).

| Sex | Patient category |  | TB type |  |
| --- | --- | --- | --- | --- |
|  | New, n (%) | Retreatment, n (%) | PTB, n (%) | EPTB, n (%) |
| Male | 83 (44.6) | 17 (9.1) | 37 (19.9) | 63 (33.9) |
| Female | 77(41.4) | 9 (4.9) | 33 (17.7) | 53 (28.5) |
| **Total** | **160 (86.0)** | **26 (14.0)** | **70 (37.6)** | **116 (62.4)** |

n: number of cases in each category; TB: Tuberculosis; PTB: Pulmonary tuberculosis; EPTB: Extra pulmonary tuberculosis

EPTB was clinically characterized on about 62.4% (116/186) TB-positive cases in the study area, and 86.0% (160/186) of them were newly diagnosed for TB (Table 2).

### Risk factor assessment for TB prevalence

Patients orign (p<0.001), age groups (p<0.001) and TB history in the family were significantly associated (p<0.001) with TB prevalence. However, sex, educational status and raw milk consumption habits were not significantly associated with the occurrence of all forms TB (p> 0.05 for all) (Table 3). Similarly, no significant association was observed between any of the different demographic or clinical characteristics and the prevalence of EPTB among the study participants (p>0.05 for all) (Table 3).

**Table 3.** Association of risk factors to EPTB in humans, South Gondar Zone, northwest Ethiopia (2015-2017).

| Characteristics | All forms TB*<br>(%) | EPTB (%) | p-value |
| --- | --- | --- | --- |
| **Age group (year)** |  |  |  |
| <18 | 21(11.3) | 17(81.0) |  |
| 18-30 | 66 (35.4) | 43(65.2) | 0.56 |
| 31-43 | 50 (26.9) | 25(50.0) | 0.23 |
| 44-56 | 37 (19.9) | 25(67.6) | 0.66 |
| >56 | 12 (6.5) | 6(50.0) | 0.41 |
| **Sex** |  |  |  |
| Male | 100 (53.8) | 63(63.0) |  |
| Female | 86 (46.2) | 53(61.6) | 1.00 |
| **Education status** |  |  |  |
| Illiterate | 117 (62.9) | 71(60.7) |  |
| Adult education | 11(5.9) | 8(72.7) | 0.71 |
| Primary school | 37(19.9) | 24(64.9) | 0.82 |
| Secondary school | 15(8.1) | 9(60.0) | 1.00 |
| Higher education | 6 (3.2) | 4(66.7) | 0.88 |
| **TB history in the family** |  |  |  |
| Yes | 52 (28.0) | 37(71.2) |  |
| No | 134 (72.0) | 79(59.0) | 0.46 |
| **Consumption of raw milk** |  |  |  |
| Yes | 133 (71.5) | 87(65.4) |  |
| No | 53 (28.5) | 29(54.7) | 0.50 |
| **Patient category** |  |  |  |
| New | 160 (86.0) | 103(64.4) |  |
| Retreatment | 26 (14.0) | 13(50.0) | 0.48 |

\*Both pulmonary and extra pulmonary TB

### RD9-Deletion typing

A 59.7% (111/186) culture positivity was obtained from the active TB cases. Of which, 59.5% (66/111) was isolated from extra pulmonary TB (EPTB) patients. The molecular typing of culture positive isolates using RD9-based PCR revealed that all isolates had intact RD9 locus and were subsequently classified as *M. tuberculosis.*

### Spoligotyping patterns of *M. tuberculosis* isolates

A total of 111 isolates which were confirmed all to be *M. tuberculosis*, by RD9-based PCR test, subjected to spoligotyping. Among these, the patterns of 96 isolates were interpretable and grouped into 35 different spoligotype patterns. From the 35 patterns, 22 patterns were shared types and consisted of 79 isolates (Table 4).

**Table 4.**
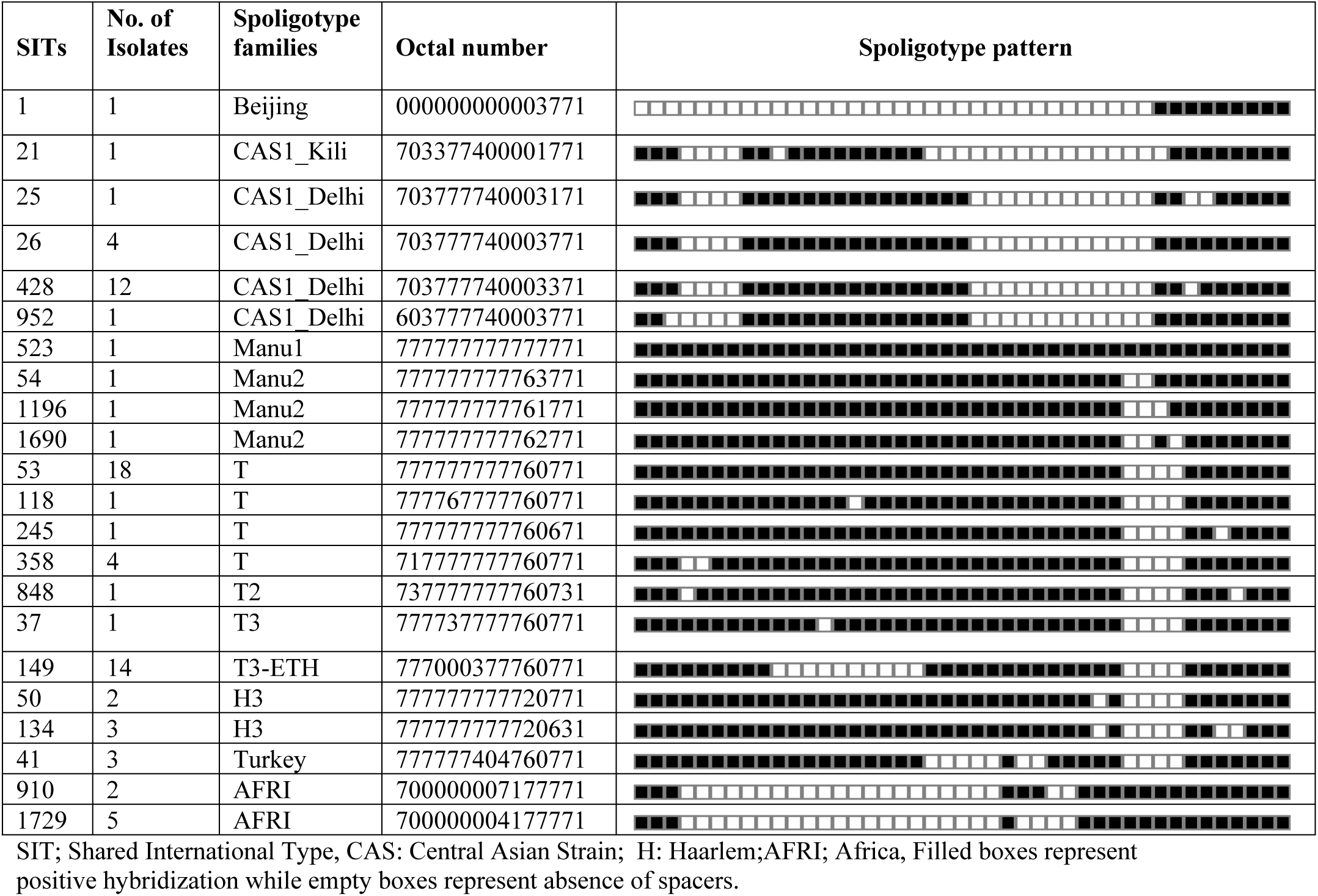
Spoligotype patterns of 22 shared types and their corresponding families identified from a total of 96 *M. tuberculosis* strains isolated in South Gondar Zone, northwest Ethiopia (2015-2017).

The dominantly identified SITs were SIT53 (Lineage 4), SIT149 (Lineage 4), and SIT428 (Lineage 3), each consisting of 18, 14 and 12 isolates, respectively (Table 4). These three SITs consisted of 45.8% of the total isolates. The remaining 13 were not registered in the database and were ‘orphans’ strains (Table 5).

**Table 5:**
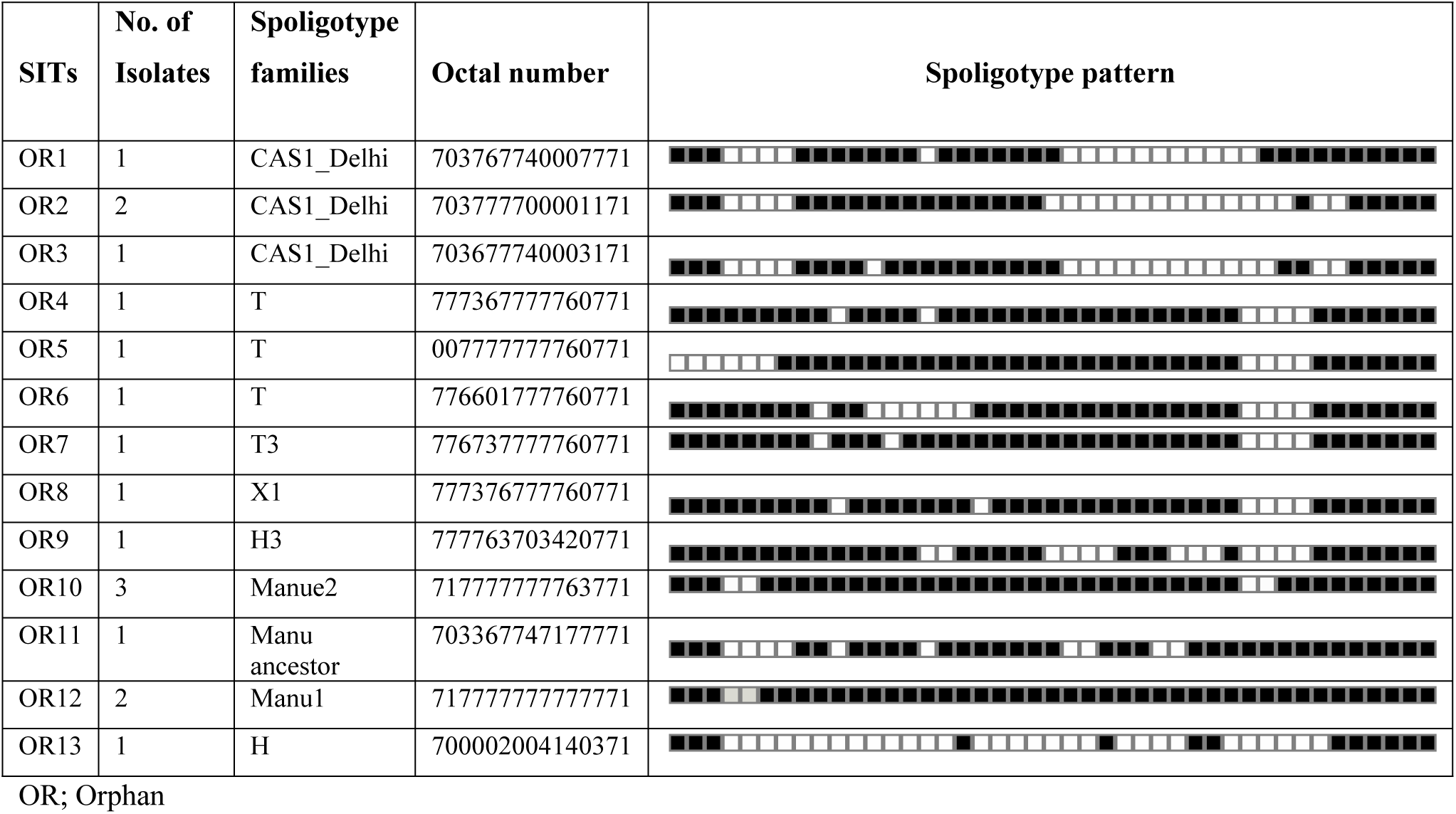
Spoligotype patterns of 13 orphan strains and their corresponding families identified from a total of 96 *M. tuberculosis* isolates collected in tuberculosis patients in South Gondar Zone, northwest Ethiopia (2015-2017).

The two dominant major lineages identified in this study were Lineage 4 (Euro-American, 62.5%) and Lineage 3 (Central Asia, 26.0%). MTBC strains identified as lineage 7 (Afri), Lineage1 (East African Indian) and Lineage 2 (Beijing) were only identified in seven, three and one patients, respectively. The three dominantly identified families were T, CAS and Manu, each consisting of 46.9%, 24.0% and 10.4% of the isolates, respectively (Tables 4 and 5).

### Association of patient characteristics with *M. tuberculosis* genotype clustering

Higher percentages of the total 96 *M. tuberculosis* isolates were identified from Dera District (30.2%) followed by Fogera District (22.9%). The clustering rates of strains isolated from these two districts were 89.7 and 72.7, respectively. The least number of *M. tuberculosis* isolates were identified from Este (3.1%) with a clustering rate of 33.3% (Table 6).

**Table 6.**
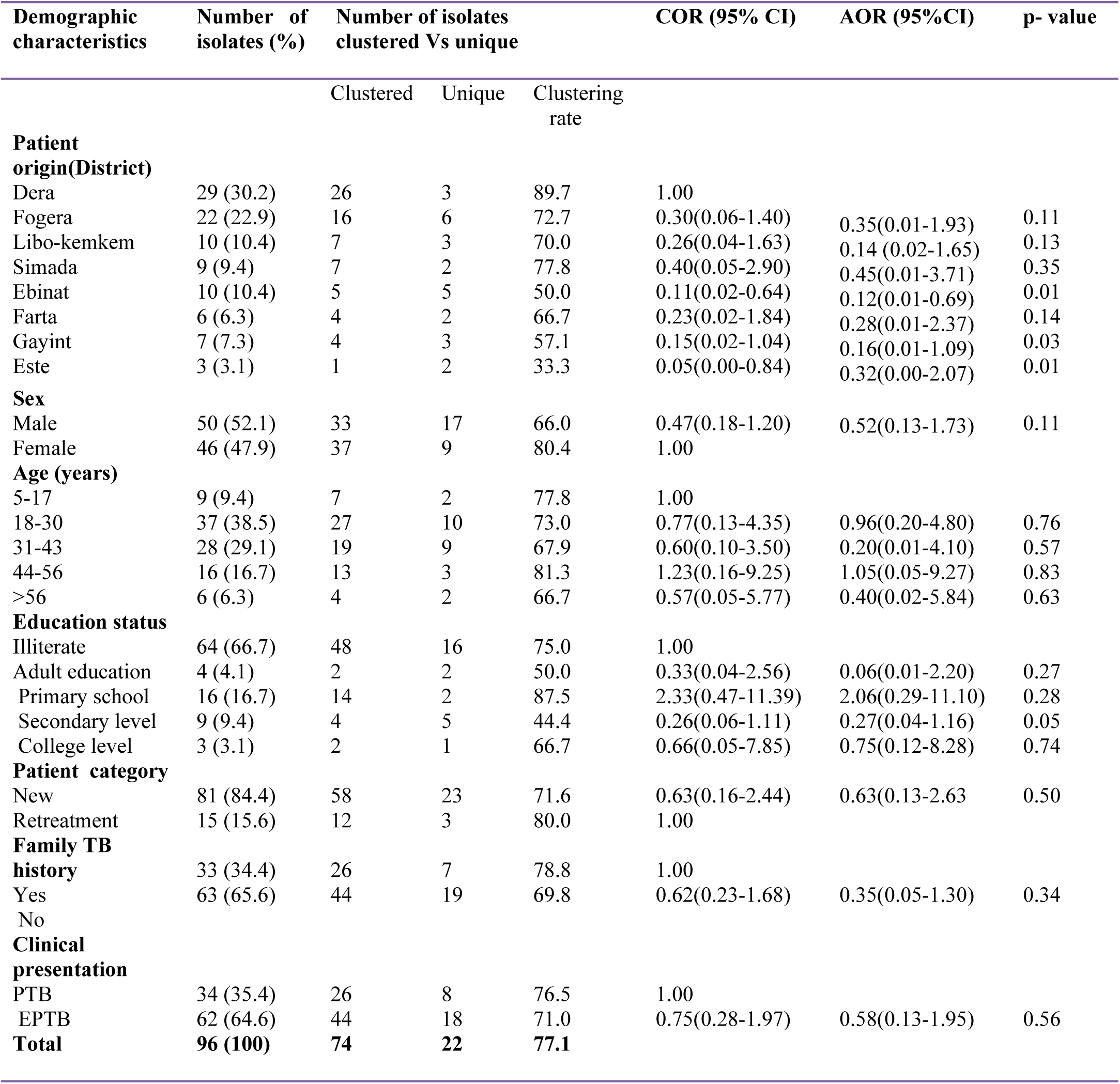
Demographic and clinical characteristics of the study participants and their association with spoligotype clustering of TB isolates from South Gondar Zone, northwest Ethiopia (2015-2017).

| Demographic characteristics | Number of isolates (%) | Number of isolates clustered Vs unique |  |  | COR (95% CI) | AOR (95%CI) | p- value |
| --- | --- | --- | --- | --- | --- | --- | --- |
|  |  | Clustered | Unique | Clustering rate |  |  |  |
| **Patient origin(District)** |  |  |  |  |  |  |  |
| Dera | 29 (30.2) | 26 | 3 | 89.7 | 1.00 |  |  |
| Fogera | 22 (22.9) | 16 | 6 | 72.7 | 0.30(0.06-1.40) | 0.35(0.01-1.93) | 0.11 |
| Libo-kemkem | 10 (10.4) | 7 | 3 | 70.0 | 0.26(0.04-1.63) | 0.14 (0.02-1.65) | 0.13 |
| Simada | 9 (9.4) | 7 | 2 | 77.8 | 0.40(0.05-2.90) | 0.45(0.01-3.71) | 0.35 |
| Ebinat | 10 (10.4) | 5 | 5 | 50.0 | 0.11(0.02-0.64) | 0.12(0.01-0.69) | 0.01 |
| Farta | 6 (6.3) | 4 | 2 | 66.7 | 0.23(0.02-1.84) | 0.28(0.01-2.37) | 0.14 |
| Gayint | 7 (7.3) | 4 | 3 | 57.1 | 0.15(0.02-1.04) | 0.16(0.01-1.09) | 0.03 |
| Este | 3 (3.1) | 1 | 2 | 33.3 | 0.05(0.00-0.84) | 0.32(0.00-2.07) | 0.01 |
| **Sex** |  |  |  |  |  |  |  |
| Male | 50 (52.1) | 33 | 17 | 66.0 | 0.47(0.18-1.20) | 0.52(0.13-1.73) | 0.11 |
| Female | 46 (47.9) | 37 | 9 | 80.4 | 1.00 |  |  |
| **Age (years)** |  |  |  |  |  |  |  |
| 5-17 | 9 (9.4) | 7 | 2 | 77.8 | 1.00 |  |  |
| 18-30 | 37 (38.5) | 27 | 10 | 73.0 | 0.77(0.13-4.35) | 0.96(0.20-4.80) | 0.76 |
| 31-43 | 28 (29.1) | 19 | 9 | 67.9 | 0.60(0.10-3.50) | 0.20(0.01-4.10) | 0.57 |
| 44-56 | 16 (16.7) | 13 | 3 | 81.3 | 1.23(0.16-9.25) | 1.05(0.05-9.27) | 0.83 |
| >56 | 6 (6.3) | 4 | 2 | 66.7 | 0.57(0.05-5.77) | 0.40(0.02-5.84) | 0.63 |
| **Education status** |  |  |  |  |  |  |  |
| Illiterate | 64 (66.7) | 48 | 16 | 75.0 | 1.00 |  |  |
| Adult education | 4 (4.1) | 2 | 2 | 50.0 | 0.33(0.04-2.56) | 0.06(0.01-2.20) | 0.27 |
| Primary school | 16 (16.7) | 14 | 2 | 87.5 | 2.33(0.47-11.39) | 2.06(0.29-11.10) | 0.28 |
| Secondary level | 9 (9.4) | 4 | 5 | 44.4 | 0.26(0.06-1.11) | 0.27(0.04-1.16) | 0.05 |
| College level | 3 (3.1) | 2 | 1 | 66.7 | 0.66(0.05-7.85) | 0.75(0.12-8.28) | 0.74 |
| **Patient category** |  |  |  |  |  |  |  |
| New | 81 (84.4) | 58 | 23 | 71.6 | 0.63(0.16-2.44) | 0.63(0.13-2.63) | 0.50 |
| Retreatment | 15 (15.6) | 12 | 3 | 80.0 | 1.00 |  |  |
| **Family TB history** |  |  |  |  |  |  |  |
| Yes | 33 (34.4) | 26 | 7 | 78.8 | 1.00 |  |  |
| No | 63 (65.6) | 44 | 19 | 69.8 | 0.62(0.23-1.68) | 0.35(0.05-1.30) | 0.34 |
| **Clinical presentation** |  |  |  |  |  |  |  |
| PTB | 34 (35.4) | 26 | 8 | 76.5 | 1.00 |  |  |
| EPTB | 62 (64.6) | 44 | 18 | 71.0 | 0.75(0.28-1.97) | 0.58(0.13-1.95) | 0.56 |
| **Total** | **96 (100)** | **74** | **22** | **77.1** |  |  |  |

Patient origin and educational background were observed to be significantly associated patient characteristics with clustering level of the strains. TB patients from Este and Ebinat districts were less likely to have clustered strains than those from Dera[(Este vs. Dera: odds ratio [AOR] =0.32; 95% CI:0.00-2.07; p =0.01)] and [(Ebinat vs. Dera: AOR= 0.12; 95% CI: 0.01-0.69;p =0.01)].

Similarly, as compared to Dera, there was lower odd of strain clustering in samples from Gayint district (Gayint vs. Dera: AOR= 0.16; 95 % CI:0.01-1.09;p =0.03). Clustering of strains was more than three times higher in sample from illiterates than from those with secondary school education level (Secondary vs. Illiterates: AOR=0.27; 95% CI: 0.04-1.16; p = 0.05). Clustering level, however, was not significantly associated with the other demographic and clinical characteristics of the study participants (Table 6).

A total of 74 (77.1%) isolates were grouped into 13 different clusters of strains which ranged from 2 to 18 isolates. The remaining 22 (22.1%) isolates had single strains (unique strains). The overall clustering rate was 77.1 (Table 6). Higher number (26) of clustered strains occurred in the strains of the Dera district (Table 6).

## Discussion

The 6.3% overall prevalence of all forms of TB recorded in individuals visiting public health institutions in the present study was comparable with the report (6.2%) from northeast Ethiopia [6], whereas relatively lower prevalence of TB was reported among private health clinic attendees (5.4%) [3] and in the prison population in Gondar, northwest Ethiopia (5.3%) [8]. A 4.6% lower prevalence was reported in general population living in southwest Ethiopia [22]. Lower TB prevalence was also reported in other African countries such as Zambia (4.3%) and Botswana (4.2%) [23, 24]. On the contrary, some studies in Ethiopia had revealed higher prevalence of TB within the range of 7.5% to 17.3% [10, 25-27]. Higher TB prevalence was also reported in South Africa (30.2%) [28]. The observed variations in TB prevalence might be due to differences in the quality of diagnostic techniques used as well as variations in the living condition of the study populations.

About 62.4% of smear positive TB cases in the present study were EPTB. This was higher than reports of other two studies conducted in northwest Ethiopia (59.8% and 28.3%) and a study from pastural communities of East Ethiopia (22.6%) [29-31]. Lower prevalence of EPTB was also reported in northern Nigeria (32.6%) and Cameroon (12,7%) [32, 33]. The reason for the high prevalence of EPTB in the present study might be due to the fact that many of the study participants were from referal hosptals where highly suspected EPTB cases diagnosed.

In this study, higher TB prevalence was found in Dera and Fogera districts. In consistent to our finding, it was reported that place of residence was found to be a risk factor for TB [6, 8]. The relative higher population size of the study sites [11], living conditions in the study area and migration of working staff to these areas from other districts in the zone might have contributed for the recorded variation in TB prevalence among the different sites in the study area.

In agreement with previous studies in other parts of Ethiopia [3, 34], age of study participants was observed to be significantly associated to TB prevalence in the present study. Specifically, individuals within the age range of 31 to 43 years were observed to be infected with TB in a higher proportion (25.6%). In South Africa, 88.2% prevalence of TB in the 31 to 35 years of age study participants was reported [7]. The reason for the observed high prevalence of the disease in these age groups might be due to their greater chance of contact to the community due to their more interactive day to day life activities. The global high prevalence of HIV/AIDS in these age groups might have also contributed for the observed high TB prevalence [35, 36].

In the present study none of the demographic or clinical characteristics of the study participants were observed to be significantly associated to the development of EPTB. Consistently, a study in Gondar reported a non-significant association of age, sex, educational status, residence and occupational status of the respondents with EPTB infection [37]. However, some studies elsewhere reported that occurence of EPTB depends on the region, ethnic background, age, underlying disease, immune status of the patient as well as genotype of the *Mycobacterium tuberculosis* strain [38-40]. Such differences might be due to differences in the use of more advanced diagnostic techniques than smear microscopy that was used in the present study.

The identification of *M. tuberculosis* as the only *Mycobacterium* species in the present study, using RD9-based PCR, was in agreement with previous reports in other parts of Ethiopia in which all or the majority of the isolates found from human TB cases were *M. tuberculosis* [41-44]. In contrast, previous studies conducted in large scale commercial farms and pastoral communities in lowland areas suggested the contribution of *M. bovis* to the overall burden of TB in humans [45, 46]. The reason for the difference in *Mycobacterium* species prevalence in this study and previous studies might be due to the low TB infection rate in cattle owned by smallholder farmers that participated in the present study.

The relatively low, 36.5% (35/96), genetic diversity of spoligotype strains in this study was consistent with similar studies conducted in Ethiopia [41-43, 47]. This might be an indication that genetic diversity of spoligotype strains is limited in Ethiopia.

The most prevalent *M. tuberculosis* strains in this study, SIT53 (lineage 4) and SIT149 (lineage 4), have been frequently reported from other parts of Ethiopia [47-50], while strain SIT428 (lineage 3) appears to be specific to the study area since it was not reported in previous studies from other parts of Ethiopia and East Africa. However, SIT428 was reported in some parts of Asia such as India [51] and Iran [52]. Despite this, no any epidemiological link was observed to the specificity of this strain in the study area.

In agreement with previous studies conducted in Ethiopia [44, 50, 53, 54], lineage 4 (Euro-American) was the dominant (62.5%) major lineage identified, while T3-ETH, among the most prevalent T family, constituted a considerable share (14.6%) in this study. This is in agreement with previous studies that reported a high proportion of T3-ETH from mycobacteria isolates from Ethiopia [42, 47, 55].

In the present study, except partly for the patients’ origin, no significant association was observed between socio-demographic factors and mycobacterial strain clustering. Similarly, studies conducted in northwest Ethiopia [56] and elsewhere [57, 58] also reported no significant association in this respect. This lack of association may be explained by high force of interaction (proportion of susceptible individuals who have become infected in a specified period), which would result in considerable rates of TB infections [58]. In this study, the observed high clustering rates of *M. tuberculosis* strains in Dera (89.7%), Simada (77.8) and Fogera (72.7%) were possibly due to the relatively high population density, suggesting an ongoing TB transmission in the study area.

The present study has some limitations in including other patient characteristics such as the HIV status of the study participants and failure in the use of more advanced molecular techniquies than spoligotyping to better support epidemiological links to the occurrence of TB in the study area.

## Conclusions

The present study revealed high prevalence of all forms of TB with a higher proportion of EPTB and high clustering rate of the *M. tuberculosis* isolates. Moreover, patients’ origin, age and family history of TB were found to be the main risk factors of TB. Hence, appropriate TB control strategies should be implemented in the study area.

## Acknowledgements

We also would like to thank all laboratory working staff and administrators at the Regional Health research Center, Bahirdar and ALIPB, Addis Ababa University, for their support in the development of this study. We also greatly thank health officials, laboratory workers and study participants in the study area, without whom this study would have not been completed.

---

**S1 Appendix: Informed consent form for the study participants.** The consent form was designed for the study participants or witnesses in case of illiterate ones who attend health facilities during the study period in South Gondar Zone, northwest Ethiopia.

